# Convergent acoustic community structure in South Asian dry and wet grassland birds

**DOI:** 10.1101/2020.08.07.241612

**Authors:** Sutirtha Lahiri, Nafisa A. Pathaw, Anand Krishnan

## Abstract

Although the study of bird acoustic communities has great potential to provide valuable conservation data, many aspects of their assembly and dynamics remain poorly understood. Grassland habitats in South Asia comprise distinct biomes with a unique avifauna, presenting an opportunity to address how community-level patterns in acoustic signal space arise. Similarity in signal space of different grassland bird communities may be due to phylogenetic similarity, or because different bird groups partition the acoustic resource, resulting in convergent distributions in signal space. Here, we quantify the composition, signal space and phylogenetic diversity of bird acoustic communities from the dry semiarid grasslands of Northwest India and the wet floodplain grasslands of Northeast India. We find that acoustic communities occupying these distinct biomes exhibit convergent signal space. However, dry grasslands exhibit higher phylogenetic diversity, and the two communities are not phylogenetically more similar than expected by chance. The Sylvioidea encompasses half the species in the wet grassland acoustic community, with an expanded signal space compared to the dry grasslands. Thus, dry and wet grassland communities are convergent in signal space despite differences in phylogenetic diversity. We therefore hypothesize that different clades colonizing grasslands partition the acoustic resource, resulting in convergent community structure across biomes. Many of the birds we recorded are highly threatened, and acoustic monitoring will support conservation measures in these imperiled, yet poorly-studied habitats.

## Introduction

The acoustic signals of different bird species may diverge to minimize competitive overlap, leading to overdispersion or uniform distribution of species within acoustic signal space. Each species within the acoustic community occupies a different region of this space (Nelson and Marler 1990; Luther 2009; Krishnan 2019). However, the role of community phylogenetic structure in driving these signal space patterns remains poorly understood. To illustrate, if two biogeographically distinct acoustic communities partition the acoustic resource, they are predicted to exhibit convergent distributions in signal space (trait space), in spite of dissimilar phylogenetic compositions. Alternatively, communities may possess similar or divergent distributions in signal space simply as a consequence of phylogenetic similarity or dissimilarity. Quantifying the species compositions of different bird acoustic communities, their respective signal (or trait) spaces, and phylogenetic similarity are necessary to test these predictions (Webb 2000; Webb et al. 2002; Cavender-Bares et al. 2004).

Grassland habitats are widespread across both tropical and temperate regions of the world, yet are highly threatened by habitat destruction (Nerlekar and Veldman 2020). Because birds are a visible and vocal component of grassland fauna, they are a valuable indicator of ecosystem health (Vickery and Herkert 1999). Most grassland birds nest on or close to the ground, and many possess highly specific habitat requirements. As a result, habitat destruction is causing steep declines in grassland bird populations (Vickery and Herkert 1999; Dutta et al. 2011; Rahmani 2012; Hill et al. 2014; Correll et al. 2019). However, until recently, grassland birds, particularly in tropical regions, have received relatively little study or conservation attention compared to forest birds (Krishnan 2019). This lacuna is particularly pronounced in the Indian Subcontinent, which possesses diverse grassland habitats (Ratnam et al. 2016) occurring along a range of rainfall regimes. These include the semiarid dry grasslands of the Thar desert in the northwest, and the wet alluvial floodplain grasslands of the Gangetic-Brahmaputra floodplains in the northeast (Dabadghao and Shankarnarayan 1973). Although occurring in different ecoregions, bird species are thought to have speciated across the boundary between dry and wet grasslands (Ripley and Beehler 1990), and their acoustic communities remain unstudied. An examination of community composition and phylogenetic similarity across these two biomes presents an opportunity to understand how acoustic communities are assembled, and also use this data as a baseline for non-invasive monitoring.

Here, we study the avian acoustic communities of dry grasslands in Northwest India, and of wet floodplain grasslands in Northeast India. First, we assess the species compositions, distributions and diversity of birds in these acoustic communities. Secondly, we quantify the signal space occupied by vocal birds in each habitat, and assess whether the two exhibit convergent community structure (i.e. distributions in signal space). Finally, we test whether community structure arises from phylogenetic similarity between the communities (similar species or close relatives in each habitat), or from different bird groups expanding to fill the same signal space (i.e. phylogenetically dissimilar communities). Our findings, some of the first detailed acoustic data from these habitats, have great value in long-term conservation monitoring of threatened grassland biomes and their unique avifauna.

## Materials and Methods

### Study sites

We conducted fieldwork in two protected areas, which contain some of the best-preserved semiarid and floodplain grasslands in India (which we refer to as “dry grassland” and “wet grassland” henceforth) (Figure 1). For wet grasslands, we carried out acoustic sampling in the D’Ering Wildlife Sanctuary in East Siang District, Arunachal Pradesh (27°51’-28°5’ N, 95°22’-95°29’ E). This sanctuary consists of seasonally inundated riverine islands, or *chaporis*, with a *Phragmites-Saccharum-Imperata* type grassland structure (Dabadghao and Shankarnarayan 1973; Rahmani 2016a). Within this habitat, two grassland types exist: tall (3-4m) grassland that does not undergo burning, and shorter (1m) grassland that is managed using annual controlled burns. The latter also contains forested patches and scattered trees such as *Bombax ceiba* (Rahmani 2016a). The grasslands are in close proximity to water, with human habitation on the other side of the Siang river. We sampled tall-grass habitat in the Borghuli range of the sanctuary, and short-grass habitat in the Anchalghat range on a different *chapori*. Our landscape-level analysis considered the overall acoustic community across these sites, as tall and short grassland often exist in close proximity in other areas of the floodplain, and therefore a landscape-level analysis is the most appropriate assessment of the overall acoustic community.

**Figure 1:**
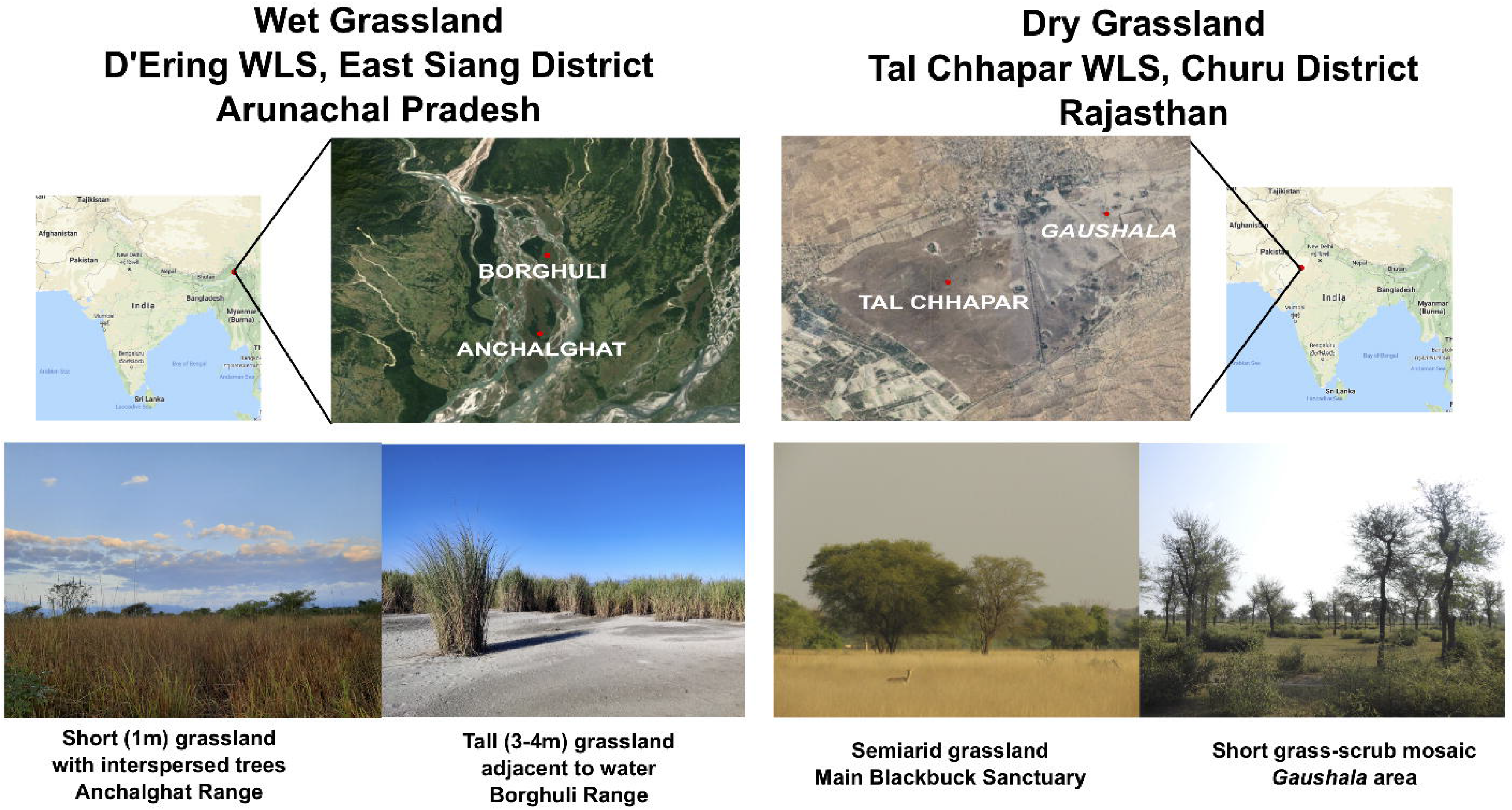
Locations of the two sanctuaries, showing the various habitats sampled. Photos courtesy Taksh Sangwan (left) and Ram Mohan (right).

For dry grasslands, we sampled in the Tal Chhapar Wildlife Sanctuary in Churu District, Rajasthan (27°47′ N, 74°26′ E). This sanctuary is located on the edge of the Thar Desert, containing xerophilous *Dichanthium-Cenchrus-Lasiurus* type grasslands, with intermixed scrub of species such as *Capparis* and *Prosopis* (Dabadghao and Shankarnarayan 1973; Kaur et al. 2020) and man-made water bodies. The village of Chhapar exists at the border of this relatively small sanctuary, and human habitation is adjacent to the grasslands. We sampled sites within the sanctuary, as well as two sites at a community grazing area (or *gaushala*) 2Km from the sanctuary boundary. This second location contains short grass habitat mixed with *Prosopis cineraria* trees, and provides additional habitat for grassland and scrub birds.

### Fieldwork and acoustic sampling

We sampled winter acoustic communities in both grassland habitats, partly to include winter migrant birds (Grimmett et al. 1998; Rasmussen and Anderton 2005) in our analysis of acoustic communities, but also because the grasslands at D’Ering WLS are inundated and inaccessible for much of the wet season (Rahmani 2016a). Therefore, the winter season is the best time to directly compare both acoustic communities at a similar time of the year. Our study at Tal Chhapar was conducted in late November-early December 2019, and our study at D’Ering shortly afterward in early January 2020.

Following previously established methodology (Krishnan 2019), we recorded the dawn singing activity of grassland habitats and used this to quantify the composition, diversity and structure of acoustic communities. Bird song was recorded using either a SongMeter SM4 (Wildlife Acoustics Inc., Maynard, MA, USA) or a Sennheiser (Wedemark, Germany) ME62 microphone connected to a Zoom (Tokyo, Japan) H6 recorder. In total, we sampled six distinct recording locations in D’Ering WLS, three in the Anchalghat range and three in the Borghuli range. At Tal Chhapar WLS, we sampled five sites, three in the main sanctuary and two in the *gaushala* (the latter were combined for site-wise analysis). Sites were selected based on the availability of suitable vegetation (extensive patches of grassland/scrub, and a safe place for the recorders). At each site, we began recording at 6AM (early dawn, just after sunrise), and recorded for at least two hours (and often over three, depending on the time taken to reach recorders and stop recording; we censused up to three hours of recording wherever available). We periodically monitored recorders owing to the risk of theft or damage by mammals.

To calculate the acoustic abundance index at the landscape level (see below), we used two hours of censused recording to ensure that each sample was equally represented, but for site-wise comparisons (also below), we used the entire censused dataset for each location. In D’Ering WLS, we sampled each site twice, resulting in a total of 36 hours of audio over 12 individual recordings. At Tal Chhapar, because there were a smaller number of sampling locations where we could place recorders, we sampled two sites 4 times each, a third 3 times and obtained two samples from the *gaushala*. This was a total of 13 individual recordings representing approximately 36 hours of audio. In one of these samples, an equipment malfunction left us with only one hour of audio, and so we excluded it from the landscape-level abundance (although we included it in site-wise analyses). Therefore, we obtained a comparable amount of audio data from both dry and wet grasslands.

### Acoustic community analysis

After dividing each recording into 10-minute segments, we identified all bird species vocalizing in each segment and constructed presence-absence matrices (one matrix for each individual recording, i.e. 12 from D’Ering and 13 from Tal Chhapar), where a value of 1 meant the species’ vocalizations were detected, and a 0 indicated absence. Two of us censused the data to minimize individual biases in detection, and also cross-verified species identity to minimize identification uncertainties. Using recordings, our observations during sampling, and online song databases, we were able to identify most vocalizations down to species. The few remaining unidentified vocalizations were almost all single detections (much less than 1% of the total) that do not, therefore, influence our community-level analysis. For further analysis, we considered species that had been detected in at least 5% of total 10-minute samples (considering the first two hours of all recordings). Next, we categorised these species (based on published information about their habitats) into grassland residents, grassland migrants and non-grassland species (both resident and migrant) (Ali and Ripley 1997; Grimmett et al. 1998; Rasmussen and Anderton 2005; del Hoyo et al. 2014). The first two categories consisted of species that regularly use grassland habitats, even if not entirely confined to them. The third category contained forest birds such as barbets and orioles, flyovers such as starling flocks and parakeets, waders and waterfowl which were detected near water, and birds from nearby human habitations, such as crows, peafowl and sparrows. Additional notes on this are in Supplementary Data. To control for any subjectivity arising from this classification, we performed analyses both on the entire acoustic community, as well as the subset of grassland species.

For each species within the two acoustic communities, we determined an “acoustic abundance index” at the landscape level, using the first two hours of each individual recording (roughly 24 hours of audio total from each community and 48 hours in total, 12 10-minute samples per matrix, across 12 matrices for each community), based on published methods (Krishnan 2019). We used the first two hours of each recording in this analysis to ensure a consistent sample size to calculate acoustic abundance; bird singing activity was typically higher in this time period. The acoustic abundance index represents the probability of detecting a species’ vocalizations if one 10-minute sample were drawn at random from each of the 12 presence-absence matrices (one for each sample). We performed this random draw 10,000 times, and took an average of the proportion of samples that contained the species (Krishnan 2019). Next, we constructed ranked-abundance curves for each habitat, and calculated Jaccard’s similarity index (i.e. shared vocal species between dry and wet grasslands), and the evenness metrics E_Q_ and E_var_, (Smith and Wilson 1996) using the codyn (Hallett et al. 2020) package in R (R Core Team 2013). For both metrics, a value of 0 implies a non-even community and 1 a perfectly even community.

Because we sampled different sites in each sanctuary, and because the number of sampling sites was smaller in the dry grasslands owing to the smaller area of Tal Chhapar (although overall sampling effort was comparable), we also compared species abundance across recording sites. This enabled us to assess how patterns at each site influenced the abundances observed at the landscape level. We calculated, for each species at each recording site, the proportion of 10 minute samples (in the total censused dataset, not just the first two hours) in which that species was detected (referred to as site-wise abundance for convenience). We carried out this analysis for six sites at D’Ering WLS and four at Tal Chhapar WLS (we pooled the two *gaushala* sites for this analysis). For visual representation, we plotted the coefficient of variation (CV) of this proportion across sites (larger CVs indicated more variance, and more heterogeneous detections across sites, indicating specificity to certain sites or habitats). To quantitatively compare how heterogeneous the distributions of vocal birds were across recording sites, we used the RAC_difference function in the codyn package to calculate the pairwise evenness difference between recording sites (Avolio et al. 2019), as well as the rank change of each species, where larger values indicate more heterogeneity between sites. We performed this analysis separately for both wet and dry grasslands.

### Signal parameter space and acoustic community structure

In order to quantify signal space of wet and dry grasslands, we calculated note parameters for each species in the acoustic community using recordings from the databases Xeno-Canto (https://www.xeno-canto.org/) and AvoCet (https://avocet.integrativebiology.natsci.msu.edu/) to ensure a good signal:noise ratio when calculating parameters. We used recordings from the same geographical region wherever possible. After labeling the notes (10-20 per species, depending on the number of notes in the recordings) in Raven Pro 1.5 (Cornell Laboratory of Ornithology, Ithaca, NY, USA), we calculated the following 8 parameters: average, maximum and minimum peak frequency, center frequency, bandwidth at 90%, average entropy, note duration, and relative time of peak (i.e. the point during the note at which the peak frequency occurred) (Krishnan 2019; Chitnis et al. 2020). Taking a species average for each of these parameters, we performed a principal components analysis on the correlation matrix using MATLAB (Mathworks Inc., Natick, MA, USA), and used the principal component scores to test for patterns within community signal space (Luther 2009). We performed three statistical analyses: first, we compared the principal component scores of dry and wet grassland acoustic communities to 100 randomly drawn uniform distributions spanning the same range of scores, using Kolmogorov-Smirnov tests (Krishnan 2019). By measuring the percentage of times each community fit to a uniform distribution (in this case, P<0.05 means they differed significantly from uniform), we tested whether they showed signatures of dispersion versus clustering in signal space, consistent with divergent acoustic signals within the community. Second, we tested whether the two communities occupied similar regions of signal space by testing whether their respective point clouds in principal component space were drawn from the same distribution, using a MANOVA with Wilk’s lambda as the test statistic on the first 3 principal components, and separately on the original parameters as well. The last test we performed built upon this previous one to also test whether the organization of points within principal component space (or community structure) was similar across the two acoustic communities. For this, we calculated the nearest-neighbour distance (NND) for species across communities (Luther 2009; Krishnan 2019). For each species in wet grassland, we calculated the Euclidean distance to its nearest neighbour in the dry grassland communities. If the two communities exhibited similar patterns of community organization in signal space, then the average NND should be lower than that of a randomly drawn community. Therefore, we constructed 10000 randomly drawn “null communities” (Chek et al. 2003) spanning the same range of principal component scores, calculated the average NND to each of these communities, and calculated the Z score of the observed NND versus this distribution of null values. All three statistical tests were performed twice, once for the entire acoustic community and once for only the grassland species.

### Phylogenetic diversity

After testing for patterns in community structure using the signal space of both acoustic communities, we tested whether both acoustic communities were phylogenetically similar. For this, we downloaded a meta-tree from the open-source Bird Tree of Life Project (https://birdtree.org/) (Jetz et al. 2012), containing 100 possible phylogenetic hypotheses for all the species within both acoustic communities. In the case of mixed-species starling flocks, we were unable to calculate separate abundance indices for each species because it was difficult to separate their vocalizations. Therefore, we included only one species from each habitat (*Acridotheres sp.*) in phylogenetic diversity analyses, although we included both in signal space calculations. When measuring phylogenetic diversity, we calculated values for each of the 100 possible trees, giving us a distribution of values. We then compared diversity metrics to each other using paired Wilcoxon signed-rank tests. Using the picante (Kembel et al. 2010) package in R, we calculated three phylogenetic alpha-diversity (within-community) indices (Faith’s phylogenetic diversity (PD), Mean Pairwise Distance (MPD) and Mean Nearest Taxon Distance (MNTD)), and two beta-diversity (between-community) indices (Community Pairwise Distance (CPD), and Community Nearest Taxon Distance (CNTD)) for each acoustic community (Tucker et al. 2017). In addition to the raw values, we also weighted phylogenetic diversity indices by the acoustic abundance index. Finally, to quantify whether the two acoustic communities exhibited congruent phylogenetic structure, we calculated the Phylogenetic Community Dissimilarity (PCD) between the two communities (Ives and Helmus 2010). Using consensus trees for both the dry and wet grassland acoustic communities, we calculated PCD by comparison to 10000 randomly reshuffled trees using the inbuilt permutation test in the pcd function of the picante package. This function outputs an overall PCD value, which is close to 1 if the two communities are as similar as expected by random chance, >1 if they are more dissimilar than expected, and <1 if they are more similar. The function additionally calculates the contribution of non-phylogenetic components (shared species, PCDc) and phylogenetic components (PCDp). This allowed us to assess whether the PCD value obtained arose because of shared species within the community versus similarities in phylogenetic structure. Again, we calculated all phylogenetic diversity metrics both for the entire acoustic community, and for grassland species.

## Results

### Avian acoustic communities of wet and dry grasslands

We identified vocalizations of 52 bird species in wet grasslands, and 68 in dry grasslands (Supplementary Data). Of these, 31 and 37 species (32 and 38 if both starling species are considered separately, see above) were recorded in >5% of 10-minute samples, and we considered these to constitute the acoustic community (see Figure 2 for examples). These species, classified according to habitat, are listed in Figure 3. Jaccard’s similarity between the dry and wet grassland acoustic communities was 0.06, as they only shared four species (*Pycnonotus cafer, Prinia inornata, Streptopelia decaocto* and *Dendrocitta vagabunda*), thus indicating large differences in community composition.

**Figure 2:**
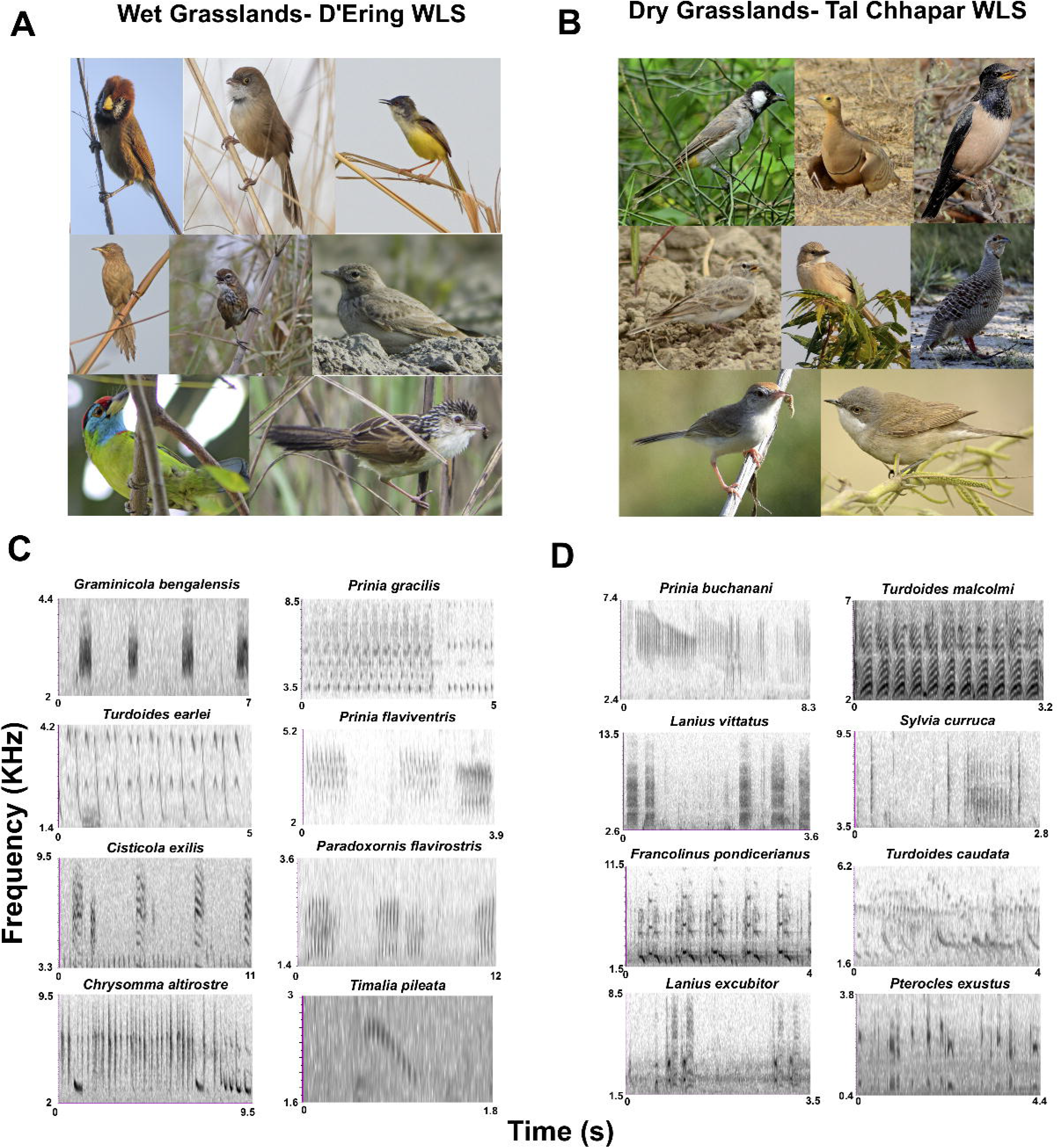
(A,B) Some of the birds comprising the acoustic community in wet (A) and dry (B) grasslands. (A) top row, left to right, photographer’s name in brackets: *Paradoxornis flavirostris* (Taksh Sangwan), *Chrysomma altirostre* (Roon Bhuyan), *Prinia flaviventris* (SL), middle row: *Turdoides earlei* (SL), *Pellorneum palustre* (Siva R), *Alaudala raytal* (SL), bottom row: *Psilopogon asiaticus* (SL), *Graminicola bengalensis* (Taksh Sangwan). (B) top row: *Pycnonotus leucotis* (SL), *Pterocles exustus* (SL), *Pastor roseus* (AK), middle row: *Calandrella brachydactyla* (SL), *Turdoides caudata* (AK), *Francolinus pondicerianus* (AK), bottom row: *Prinia buchanani* (AK), *Sylvia curruca* (SL). (C,D) Spectrograms of sample vocalizations recorded during data collection in wet (C) and dry (D) grasslands.

**Figure 3:**
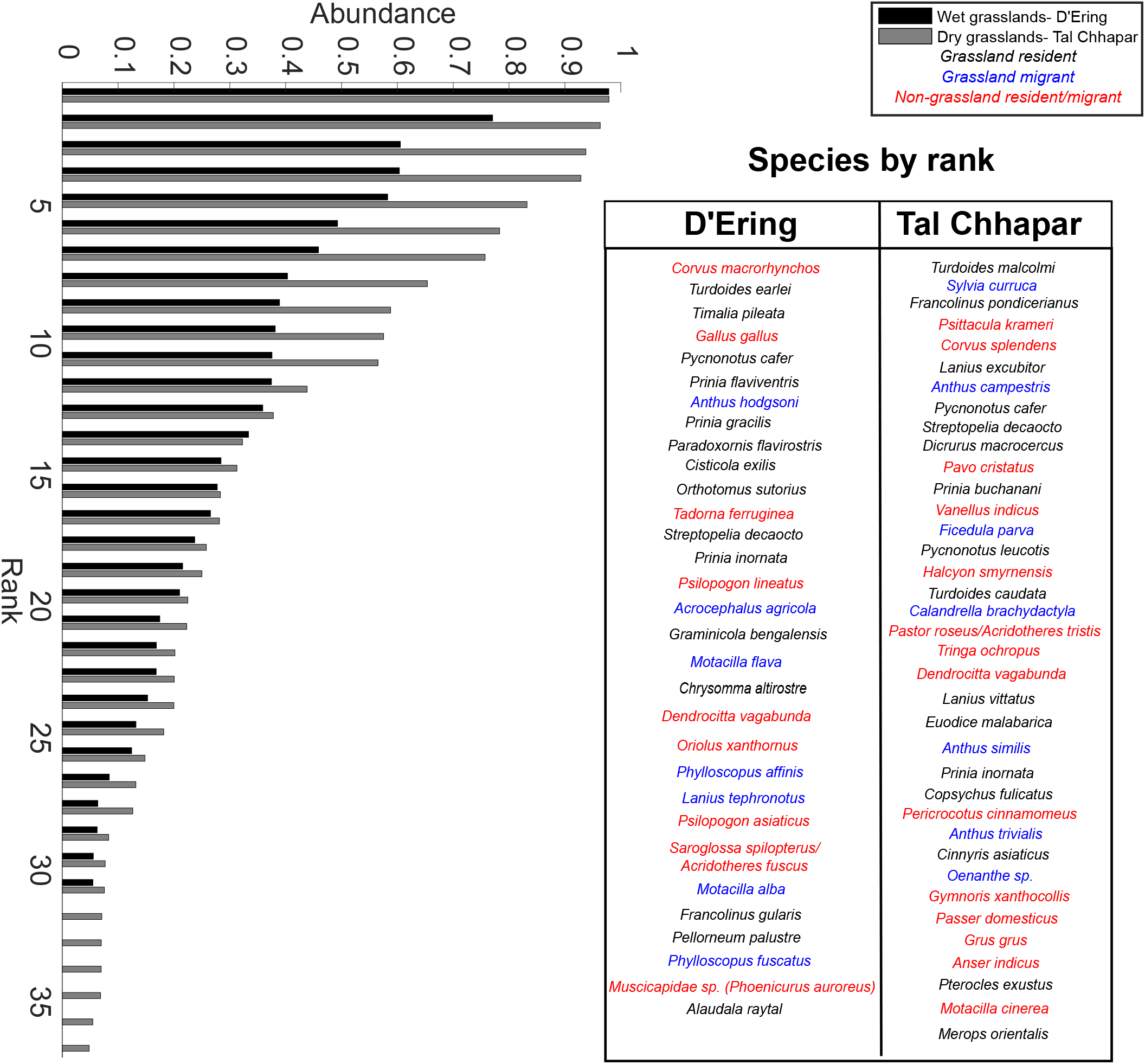
Ranked abundance distributions for both acoustic communities using the acoustic abundance index. The box contains species names in order of abundance, and color-coded by habitat preference.

Ranked-abundance distributions (Figure 3) demonstrated that higher-ranked species in dry-grassland habitats possessed acoustic abundance indices (across all sites) above 0.75, whereas only two species in wet grasslands had abundance indices above 0.6. Evenness metrics were slightly higher for wet grasslands (after normalizing for overall species diversity). This suggests that the higher abundance of vocally common species in the dry grasslands resulted a steeper drop-off of the abundance distribution, and thus slightly lower evenness across the acoustic community (E_Q_: wet grasslands 0.22, dry grasslands 0.19, E_var_: wet grasslands 0.66, dry grasslands 0.55).

### Heterogeneity across habitats in grassland bird acoustic communities

In order to investigate spatial heterogeneity in the acoustic community across recording sites and habitats, we quantified site-wise abundances for each species in both dry and wet grasslands. Site-wise abundances possessed generally higher coefficients of variation in wet grasslands than in dry grasslands, indicating more spatial heterogeneity in the wet grassland acoustic community than in the dry grasslands (Figure 4A-B). However, in general, many species exhibited heterogeneous distributions across both grasslands, as can be seen by a detailed comparison.

**Figure 4:**
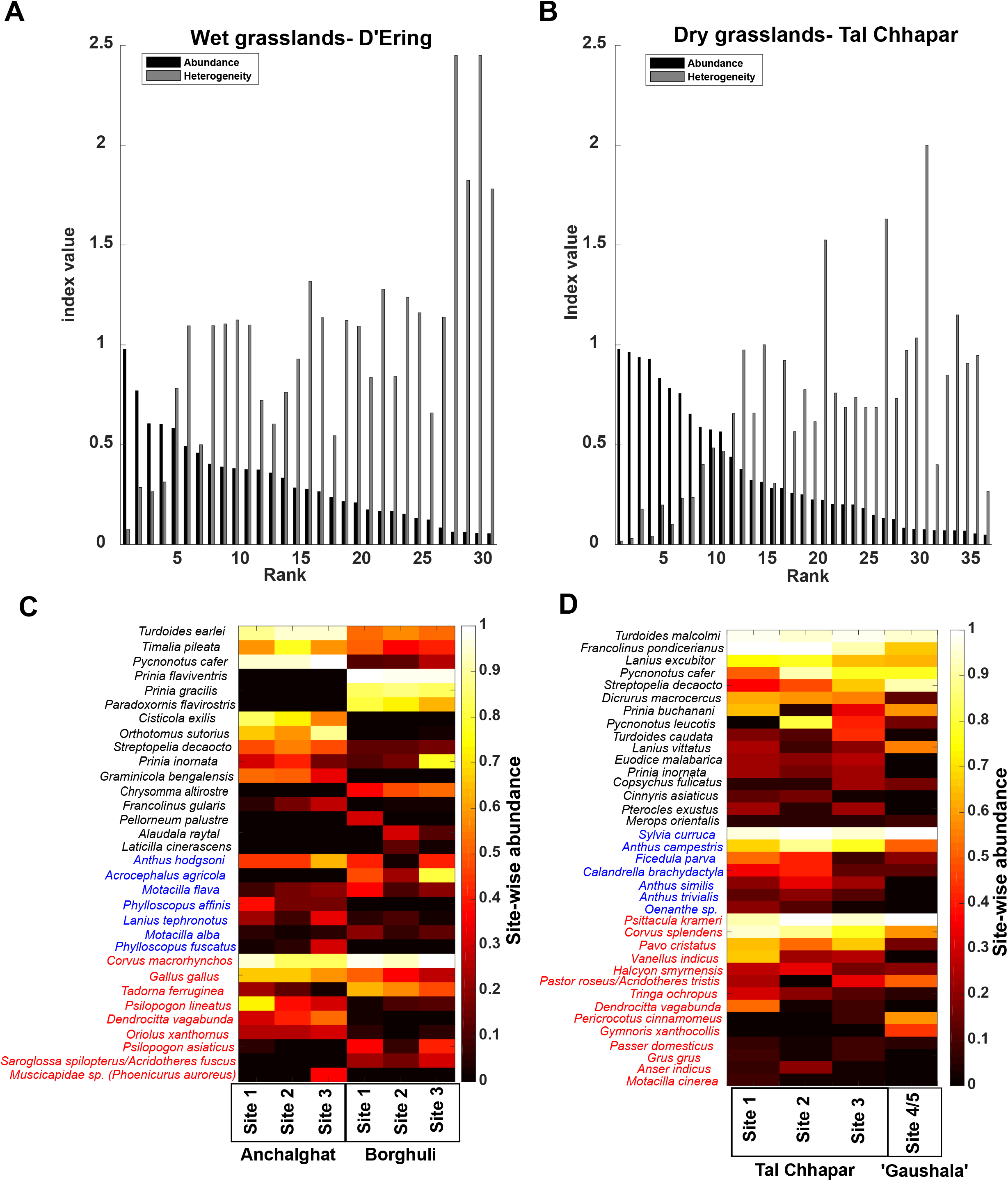
(A,B) Ranked abundance distributions (black bars) with CV of site-wise abundances for each species (grey bars). A higher CV indicates more heterogeneous distributions. (C,D) Species-wise (rows) values of site-wise abundance across recording sites (columns), with warmer colors indicating higher abundance. A checkerboard pattern is indicative of more heterogeneity in species distributions across recording sites. Species are color-coded according to their habitat preferences; note in particular the species in black (grassland residents), for whom patterns are discussed in the text. *Laticilla cinerascens* is included here solely for conservation interest on account of its rarity.

In the wet grasslands of D’Ering WLS, multiple grassland species (for example *Prinia flaviventris, Prinia gracilis, Paradoxornis flavirostris, Chrysomma altirostre* and *Alaudala raytal*) were recorded only within tall grassland. Others (*Cisticola exilis, Graminicola bengalensis, Orthotomus sutorius* and *Francolinus gularis*) were recorded either entirely or almost entirely in short grassland, and species such as *Turdoides earlei* and *Timalia pileata*, although recorded frequently at all sites, were more frequent in short grassland (Figure 4C). In the dry grasslands of Tal Chhapar WLS, some grassland species (*Turdoides malcolmi, Francolinus pondicerianus, Sylvia curruca, Streptopelia decaocto, Lanius excubitor* and *Pycnonotus cafer*) were more or less homogeneously distributed, whereas others (such as *Lanius vittatus*, *Prinia buchanani, Turdoides caudata, Pterocles exustus* and *Pycnonotus leucotis*) were more heterogeneously distributed across the landscape, with different detection levels across recording sites (Figure 4D).

In general, although pairwise evenness differences between each recording site were comparable across dry (average of the absolute evenness difference= 0.05) and wet (0.03) grasslands, rank difference (a measure of change in community composition) (Avolio et al. 2019) was higher on average for the wet grasslands (0.25, compared to 0.17 for dry grasslands), with the three tall grassland sites differing in composition from the short grassland ones (full pairwise comparisons in Supplementary Data). The dry grassland sites also differed little from each other in species composition (Supplementary Data), suggesting that the lower heterogeneity in these habitats is genuine, and not a result of sampling fewer sites.

### Wet and dry grasslands exhibit convergent acoustic community structure

Next, we compared the community signal space of wet and dry grassland acoustic communities. The first three principal components (PCs) accounted for about 85% of total variation (Supplementary Data), loading positively on frequency parameters (PC1), average entropy and bandwidth (PC2), and relative time of peak (PC3). Both communities exhibited a dispersed community structure, although the wet grassland community fit 100 randomized uniform distributions better than the dry grassland community (Kolmogorov-Smirnov tests against 100 randomized uniform distributions: wet grasslands, all species= 99%, grassland species=94%, dry grasslands, all species= 72%, grassland species= 64%; percentages indicate number of times out of a 100 where the observed distribution was similar to a uniform distribution at P greater than 0.05) (Supplementary Data) (Krishnan 2019). This dispersion is consistent with divergent signals and low interspecific overlap. Further, MANOVA on both the total acoustic community (Figure 5A) and the grassland species alone (Figure 5B) showed that the two acoustic communities occupied the same regions of signal space (all species: dF within groups=68, dF between groups=1, total dF=69, Wilk’s lambda on PCs= 0.967, P=0.53, on song parameters: lambda= 0.82, P=0.13; grassland species: dF within groups=43, dF between groups=1, total dF=44, Wilk’s lambda on PCs= 0.99, P=0.93, on song parameters: lambda= 0.81, P=0.41). This suggests that the slightly higher clustering observed in the dry grassland community is because of higher species diversity within the same region of signal space. Both communities accumulate species at broadly similar rates with distance from the origin of PC space, further supporting this assertion (Supplementary Figure 1).

**Figure 5:**
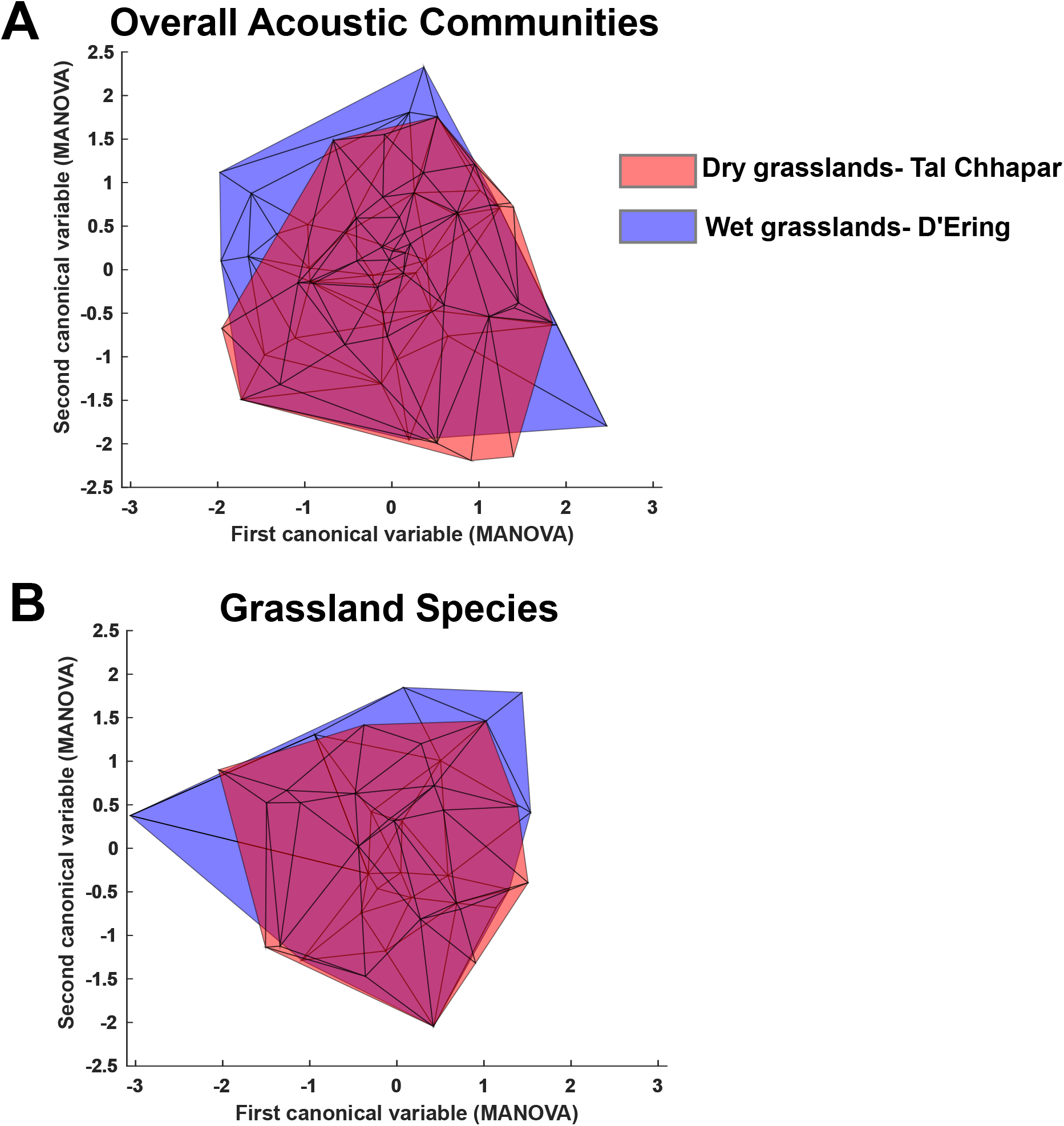
Biplot of the first two canonical variables from a MANOVA on the principal components of acoustic parameters. These parameters were measured for each species in the two acoustic communities to derive a signal space, plotted here as a minimum convex polygon using MATLAB. Note the overlap in polygons both when considering all species (A) and only grassland species (B).

Finally, a comparison of the observed between-community NNDs to those obtained from 10000 randomized null communities showed that the NNDs between dry and wet grassland acoustic communities were significantly lower than expected by chance (Z=-3.43, P<0.01). This result held true even if we only considered grassland species (Z=-3.9, P<0.01), suggesting that both these grassland biomes, in spite of possessing very few species in common, exhibit a convergent community structure in acoustic signal space.

### Dry grassland acoustic communities are higher in phylogenetic diversity

Summarizing our results so far, dry grassland acoustic communities exhibit slightly higher species diversity, but convergent community structure with wet grassland acoustic communities. We next calculated measures of phylogenetic diversity (Figure 6), to understand whether similarity in community structure was due to phylogenetic similarity of the acoustic communities. Three measures of alpha diversity, PD, MPD, and MNTD, were all significantly higher for dry grasslands than wet grasslands (Wilcoxon signed-rank test, N=100; W=0, P<0.001 for all three), even after weighting for abundance (Figure 6B-C). The same pattern held when considering only grassland species (Wilcoxon signed-rank test, N=100; W=0, P<0.001 for all three) (Figure 6E-F). Phylogenetic beta-diversity (CPD and CNTD) metrics were of roughly the same magnitude (or slightly lower, see Supplementary Data) as the corresponding alpha diversity metrics (Figure 6D,G). Weighting for abundance significantly lowered CPD but increased CNTD (Wilcoxon signed-rank test of raw versus weighted values, N=100; all species: CPD: W= 4973, CNTD: W= 1504, P<0.001, grassland species: CPD: W= 5050, CNTD: W= 0, P<0.001). Overall, though, between-community phylogenetic distances remained similar to within-community distances. A PCD value of 1.71 (all species) and 1.69 (grassland species) indicated greater dissimilarity between the communities than expected by chance. However, the PCDc value (1.87 in both cases) suggested that this difference was driven largely by differences in species composition, as PCDp (the phylogenetic component) was 0.92 (all species) and 0.91 (grassland species) (Supplementary Data). Thus, the acoustic communities were neither phylogenetically more similar nor more dissimilar than expected by random chance (Ives and Helmus 2010).

**Figure 6:**
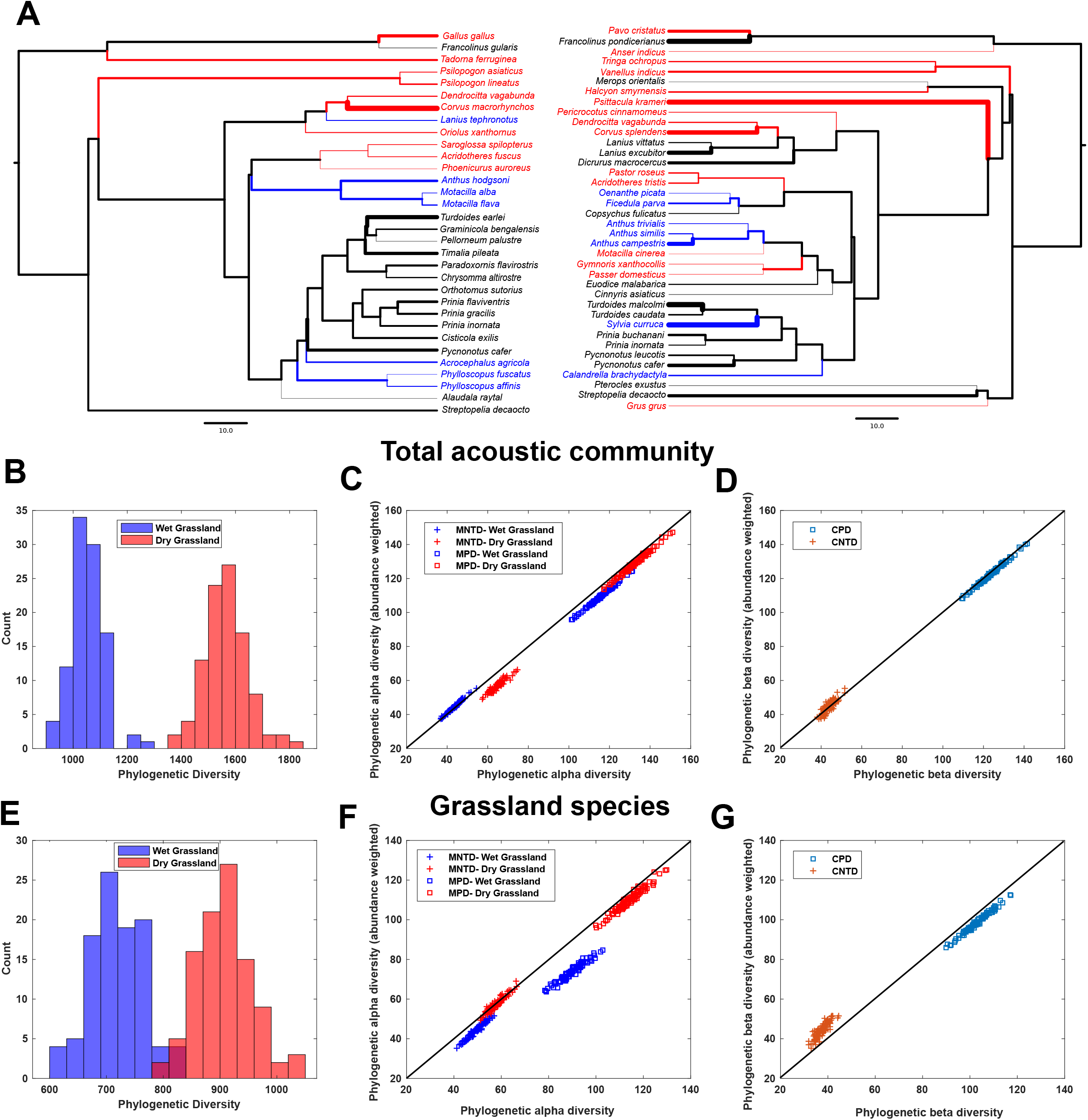
(A) Majority-rule consensus trees representing the phylogenetic relationships of species in both acoustic communities, color-coded by habitat preference as in earlier figures. Thickness of terminal branches is in proportion to abundance index for each species. (B-G) Phylogenetic alpha diversity (B,C,E,F) and beta diversity (D,G) metrics for both acoustic communities, both for all species (B-D) and grassland species (E-G). In C,D,F and G, indices are also calculated after weighting for abundance. In all scenarios, dry grasslands possess higher phylogenetic diversity.

Higher phylogenetic diversity in dry grasslands, together with a lack of phylogenetic similarity suggests that phylogenetic considerations alone do not explain convergent community structure. Because these acoustic communities are largely composed of passerine birds, we considered all non-passerines as one group, and the major passerine clades represented in our dataset (Corvoidea, Muscicapoidea, Passeroidea and Sylvioidea) as the other four groups. The dry grassland acoustic community contained 11, 6, 5, 8 and 8 species from each of these five groups respectively. However, the wet grassland acoustic community contained 6, 4, 3, 3, and 16 species respectively (considering the two starling species separately in each). The Sylvioidea (babblers, warblers, bulbuls and larks) (Alström et al. 2006) accounted for half the species in the wet grassland acoustic community, with a concomitant expansion in signal space compared to their dry grassland counterparts (Figure 7). This expansion of the Sylvioidea in wet grasslands appears to drive convergent signal space and community structure, in spite of the lower phylogenetic diversity in wet grasslands.

**Figure 7:**
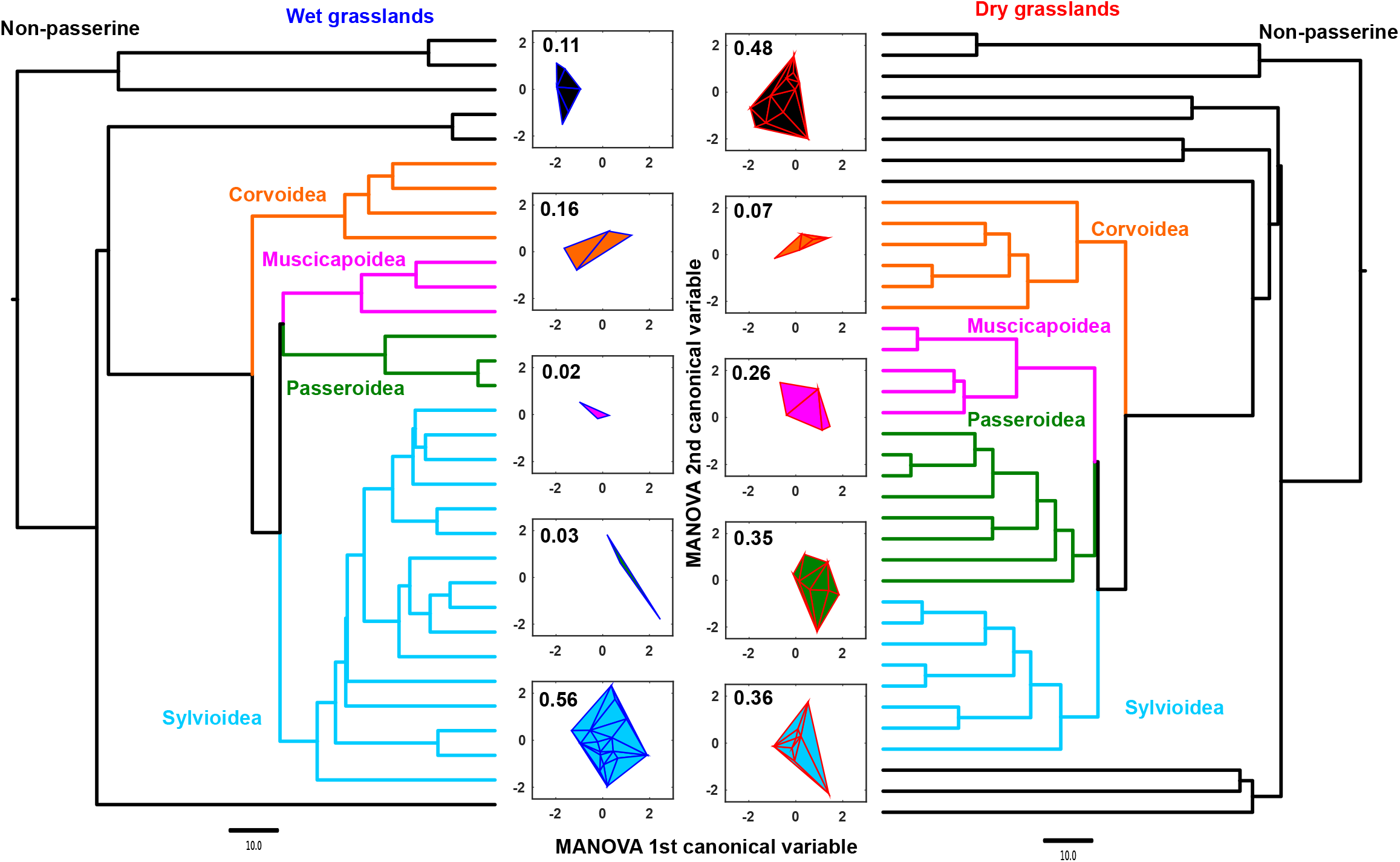
Consensus trees for each acoustic community with the major clades color-coded, and their respective signal spaces represented by minimum convex polygons in the centre. Wet grasslands are on the left (blue outline on polygons), dry grasslands on the right (red outline), and the polygons are colored according to the clade they represent. The numbers above each polygon represent its area divided by the area of the minimum convex polygon containing all species (see Figure 5). Note the dominance of the Sylvioidea in wet grasslands.

## Discussion

To summarize, we recorded diverse acoustic communities from both wet and dry grasslands in India. Although the two communities were not significantly phylogenetically similar, both exhibited convergent structure in acoustic signal space. We suggest that this is because the Sylvioidea have expanded in wet grassland signal space to occupy the same acoustic resource as the dry grassland community, even though overall phylogenetic diversity is lower in wet grasslands.

### Community bioacoustics and conservation in South Asian grasslands

Because of the multitude of threats facing tropical and subtropical grasslands in India (Ratnam et al. 2016), and the generally poor status of knowledge about their birds, bioacoustics studies have great potential to illuminate their natural history and conservation status (Blumstein et al. 2011; Campos-Cerqueira and Aide 2016; Raynor et al. 2017; Sugai et al. 2019). This is particularly true of the wet grasslands of the Brahmaputra floodplains, which have suffered extensive conversion to agriculture (Rahmani 2016b; Jha et al. 2018). Many threatened birds in this landscape remain very poorly studied. We recorded multiple globally threatened species (*Laticilla cinerascens*, *Paradoxornis flavirostris, Chrysomma altirostre, Pellorneum palustre, Francolinus gularis* and *Graminicola bengalensis*) (Rahmani 2012). Protected areas such as Tal Chhapar and D’Ering sanctuaries represent strongholds for many grassland bird species.

Grassland bird communities are often spatially heterogeneous (Vickery and Herkert 1999; Hamer et al. 2006; Hovick et al. 2014, 2015; Jacoboski et al. 2017; Londe et al. 2019), and a mix of habitats sustains the greatest diversity of birds. Broadly, our study confirms this pattern in both dry and wet grasslands in India, with more heterogeneity across recording sites in wet grasslands. However, some dry grassland species, particularly those partial to scrub cover (Ali and Ripley 1997; Rasmussen and Anderton 2005), also exhibit different abundances across recording sites. D’Ering WLS possesses distinct tall and short grasslands, compared to Tal Chhapar WLS where both grass and scrub are interspersed across sites. The dominance of the Sylvioidea in the wet grassland acoustic community, many of which are specific to certain grassland types, may also be a factor in these differences. These heterogeneous distributions are likely responsible for the slightly higher evenness in the wet grassland acoustic community, as bird species specific to either tall or short grass have similar abundances at the landscape level. For example, we recorded *Graminicola bengalensis* at all three short grassland sites, and *Chrysomma altirostre* at all three tall grassland sites, and thus their abundance indices at the landscape level were similar. Although our study primarily focused on the acoustic community at the landscape level and not on habitat variables, acoustic community structure arising from local habitat heterogeneities is an interesting future study. Our baseline data demonstrate that a community-level methodology can uncover spatial patterns within grasslands. We thus highlight the need for detailed spatial studies of bird acoustic communities in India using an occupancy framework (Dorazio et al. 2010; Furnas and Callas 2015; Campos-Cerqueira and Aide 2016; Iknayan and Beissinger 2018; Wood et al. 2019; Abrahams and Geary 2020), both to clarify habitat requirements and to study smaller-scale spatial organization in the acoustic community.

### Acoustic community structure and partitioning of the acoustic resource

Although phylogenetic diversity is higher in dry grasslands, acoustic community structure is convergent with wet grasslands, both communities exhibiting overdispersion in signal space. This overdispersion is somewhat less pronounced in dry grasslands, likely because a slightly higher number of species are occupying the same space. However, inter-community NNDs are lower than expected by chance, which is indicative of each species having a counterpart in the signal space of the other community. This may occur because grasslands filter the kinds of birds that occur in them, thus resulting in closely related species replacing each other across habitats. Their signals may resemble each other owing to their shared ancestry, thus leading to convergence in signal space (Cardoso and Price 2010; Tobias et al. 2014).

However, our phylogenetic analyses suggest the converse. Firstly, dry grassland acoustic communities have higher phylogenetic diversity, and second, the two grassland communities are no more or less phylogenetically similar than expected by chance (PCDp close to 1). The results of PCD hold even when considering all recorded species (and not just those recorded often enough to form the acoustic community, PCDp=0.9, see Supplementary data), so our results are robust to the definition of the acoustic community that we employ here. This suggests that although some close relatives may replace each other between habitats (further supported by the values of between-community CPD and CNTD being within the same range as the within-community MPD and MNTD), convergence is also responsible for the observed similarity in acoustic community structure. Some species are replaced by congeners between dry and wet grasslands (eg. *Francolinus, Turdoides*) (Ripley and Beehler 1990; Ali and Ripley 1997; Rasmussen and Anderton 2005). However, others are unique to each type of grassland. For example, *Pterocles* is found only in semiarid habitats, and *Paradoxornis* only in wet grasslands (Ali and Ripley 1997; Grimmett et al. 1998; Rasmussen and Anderton 2005; Rahmani 2012). The fact that acoustic community structure is similar in spite of these phylogenetic differences is consistent with convergence in signal space. A similar pattern holds even when considering just grassland species, indicating that the presence of villages or wooded patches does not change patterns in community structure.

The Sylvioidea (babblers, warblers, bulbuls and larks in our dataset) comprises half of the wet grassland acoustic community. This overrepresentation compared to the dry grasslands is accompanied by an increase in the signal space occupied by this clade (Figure 7), such that the overall signal space of the two communities is convergent. The Eastern Himalayas and their foothills possess the highest babbler diversity in the Indian Subcontinent (Srinivasan et al. 2014). This biogeography may partly explain why the acoustic community of the wet grasslands (occupying this region) is dominated by this clade. The expansion of the Sylvioidea’s signal space, resulting in convergent community structure with Northwestern dry grasslands, is a compelling indicator of acoustic resource partitioning. This is further supported by our analyses suggesting that species within each community are dispersed across acoustic signal space (Figure 5). Thus, we hypothesize that grassland bird acoustic communities assemble by occupying the available acoustic signal space (analogous to filling available niches) (Price et al. 2014), resulting in convergent community structure across different grassland habitats. This pattern may arise to minimize acoustic interference between relatives (Schmidt et al. 2013), or as an indirect outcome of morphological divergence between coexisting species (Krishnan and Tamma 2016). The drivers of acoustic community assembly are thus a compelling subject for future study. Quantifying community-level signal space is a valuable way to establish patterns using passive acoustic data. Thus, we suggest combining these methods with phylogenetic analyses to study spatiotemporal change in global biodiversity.

#### Acknowledgments

We are deeply indebted to the forest departments and chief wildlife wardens of Arunachal Pradesh (Letter number CWL/G/173/2018-19/Pt.VII/1585-86) and Rajasthan (Letter number F19() permission/cwlw/2017/1598) for permission and support to carry out acoustic sampling. For Arunachal Pradesh, we thank Mr. Tasang Taga, DFO, D’Ering WLS, Mr. Maksam Tayeng, Arunachal State Biodiversity Board, Taksh Sangwan and the range forest officers and forest guards of Borghuli and Anchalghat for logistical support, hospitality and accompanying us during sampling for safety. For Rajasthan, we thank Ram Mohan, Mr. Yogendra Rathod, the ACF’s office and the staff at Tal Chhapar WLS for hospitality and logistical support. We also thank Savithri Singh and Anand Prasad for assistance during sampling, Ram Mohan, Taksh Sangwan, Roon Bhuyan and Siva R for permission to use photos, Ramana Athreya and Uma Ramakrishnan for logistical advice, Viral Joshi for help with identifying bird calls, Krishnapriya Tamma and members of our research group for feedback.

#### Funding

AK is funded by an INSPIRE Faculty Award from the Department of Science and Technology and an Early Career Research Award (ECR/2017/001527) from the Science and Engineering Board (SERB), Government of India. SL received funding for this project from an Oriental Bird Club Conservation Grant, equipment support from Idea Wild, and a Rufford Foundation Small Grant (awarded to Ram Mohan, in collaboration with SL who sampled birds).

