## Supplementary Figure 1 for "Convergent acoustic community structure in South Asian dry and wet grassland birds"

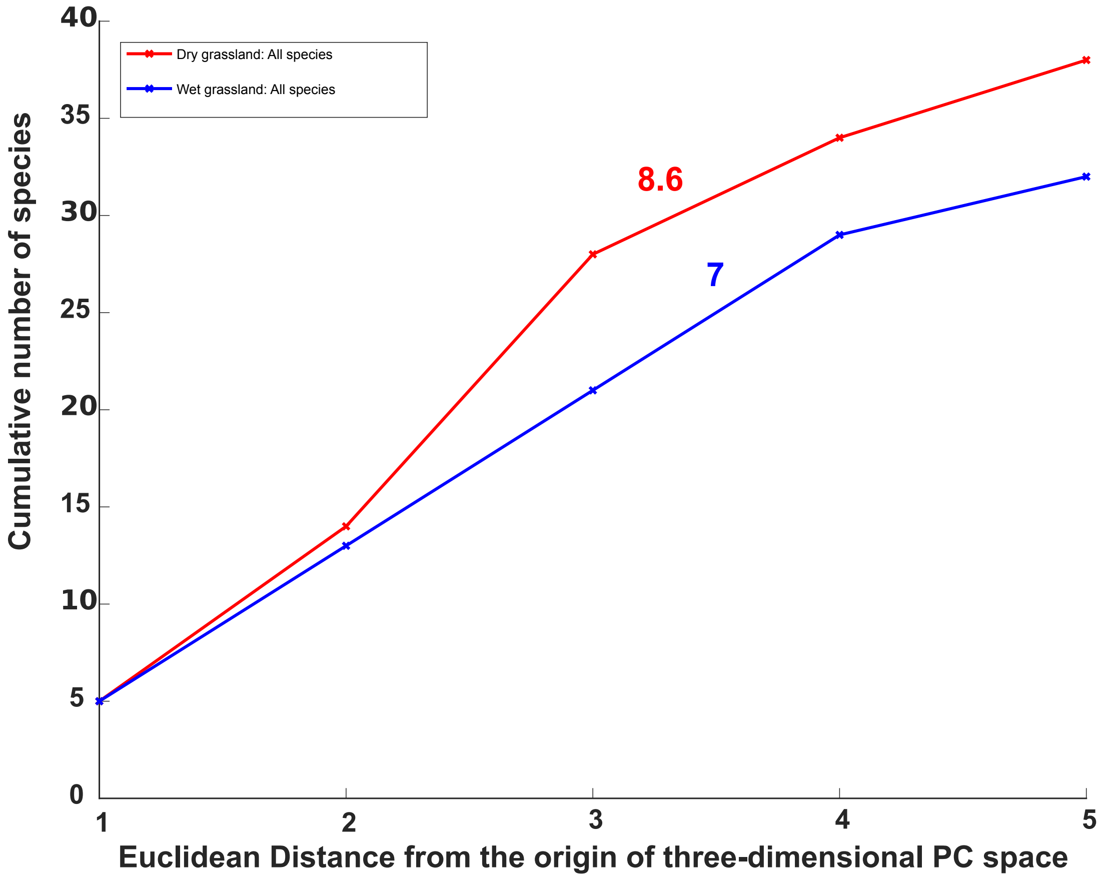


**Supplementary Figure 1: Accumulation of species with distance from the centroid of signal space. The numbers indicate the slopes obtained from a linear fit. Both communities exhibit similar distributions of species, with the main difference coming from the slightly higher number of species in dry grasslands.**
